# Seaweed Amino Acid and L-Amino Acid Improve Coriander Growth

**DOI:** 10.1101/2024.03.18.585645

**Authors:** Xingqiang Chen, Zheng Shang, Haidong Chen, Shulin Wan

## Abstract

This study investigates the impact of Seaweed amino acid (SG) and L-amino acid (LG) treatments on the growth and root development of coriander plants compared to a control group (CG). The results from Figure 1 illustrate a significant increase in biomass and foliage density for the SG and LG groups, suggesting an enhanced nutritional uptake resulting from these amino acid treatments. Both SG and LG treatments produced more vigorous growth and higher plant height compared to the CG, which received only water. Additionally, a closer inspection of coriander root systems in Figure 2 reveals an improvement in root biomass and architecture, indicating that both SG and LG applications contribute positively to root development, potentially enhancing plant resilience and yield. While both treatments showed comparable effects on root morphology, further research is required to determine if one has superior long-term benefits over the other. The findings point towards the efficacy of using amino acid treatments as bio-stimulants in agricultural practices to improve crop yield, especially in challenging growth conditions such as those found in Guangzhou, China.

## Introduction

Seaweeds, known for their biostimulant properties^1–3^, have garnered attention in sustainable agriculture for their role in enhancing plant growth and stress resistance. Coriander (*Coriandrum sativum*), a widely used culinary herb, faces cultivation challenges that demand innovative approaches to improve its yield^4^ and resilience. The versatility of Seaweed as a plant growth promoter is of particular interest in the context of coriander cultivation^5^, where stress conditions can severely impact both yield and quality.

The influence of biostimulants like Seaweed on plant growth has been increasingly documented, with studies revealing their capacity to improve nutrient uptake, enhance photosynthetic activity, and induce systemic resistance to environmental stresses^6–8^. Such multifaceted benefits make Seaweed a compelling subject for investigation within the realm of coriander cultivation^9^. This study aims to bridge the knowledge gap regarding the efficacy of Seaweed and amino acid applied through different modalities^10^, specifically, foliar application versus root drenching, and their subsequent impact on the anti-stress ability of coriander plants.

Left-handed amino acids, also known as L-amino acids, play several critical and unique roles in plants, despite being less common than their L-counterparts in nature. In plants, L-amino acids have been implicated in various physiological and biochemical processes^11, 12^. They contribute to the regulation of growth and development, acting as signaling molecules that can influence plant hormone activity and gene expression. D-amino acids are also involved in the defense mechanisms of plants, providing resistance against pathogens and pests by disrupting the biological functions of these invaders, potentially due to the inability of the invaders’ enzymes to recognize or degrade D-amino acids effectively. Additionally, they may play a role in the chiral selective absorption and metabolism of nutrients, enhancing the efficiency of nutrient uptake and utilization. Through these functions, L-amino acids are integral to the complex web of interactions and processes that support plant health, growth, and survival in a diverse range of environments.

In agricultural practice, the method of application can significantly influence the uptake and effectiveness of biostimulants. Root drenching ensures direct delivery to the root system. This research, therefore, focuses on comparing these two common application strategies for Seaweed amino acid treatment on coriander plants.

Coriander’s response to Seaweed is not solely measured in terms of visible growth parameters but also through its ability to withstand stress^13^, which is pivotal for crop survival and productivity. As environmental variables continue to be unpredictable, understanding how Seaweed affects the anti-stress ability of plants is critical. By investigating the physiological and biochemical responses of coriander to Seaweed treatments, this study endeavors to quantify the role of Seaweed in enhancing plant stress tolerance.

Finally, this introduction leads to the hypothesis that Seaweed treatments, irrespective of their application methods, would confer increased stress tolerance in coriander cultivation. This study systematically explores this hypothesis by cultivating coriander seedlings in a controlled environment and subjecting them to designated Seaweed treatments. The comprehensive analysis spans growth parameters, stress markers, and yield quality to provide a holistic understanding of the impacts of Seaweed on coriander cultivation. The outcomes are expected to offer valuable insights into optimal Seaweed application methods for enhancing the growth and stress resilience of coriander, potentially informing future horticultural practices.

## Methods

### Experimental Design and Plant Treatment

To assess the influence of amino acid applications on the growth of coriander, an experimental study was conducted in Guangzhou, China. Coriander plants were evenly distributed into three groups: one treated with Seaweed amino acid (SG), another with L-amino acid (LG), and a control group (CG). Both the SG and LG groups were subjected to a root application of their respective amino acid solutions at a concentration of 100 parts per million (ppm). This treatment was chosen to ensure that the plants had direct root access to the amino acids, potentially enhancing the uptake efficiency. The control group was irrigated solely with water, without any addition of amino acids. All groups were grown under the same environmental conditions with equal access to sunlight and water, apart from the amino acid treatment, to maintain the integrity of the comparative analysis.

### Methodology for Evaluating the Impact of Amino Acid Treatments on Coriander Root Development

The methodological approach for this study involved a controlled experimental setup to assess the effects of Seaweed amino acid (SG) and L-amino acid (LG) on coriander root development. Coriander plants were grown in Guangzhou, China, and subjected to three treatment regimens: a control group (CG) receiving only water, an SG group receiving Seaweed amino acid, and an LG group receiving L-amino acid, each at 100 ppm concentration through root application. Post-harvest, the root systems were carefully excavated and washed to remove all soil particles for an unbiased comparison. The roots were then arranged by treatment group and photographed on a uniform blue background to enhance visual contrast and facilitate the evaluation of root architecture differences among the groups. This systematic approach allowed for a direct comparison of root morphologies as influenced by the different amino acid treatments, providing insights into the potential benefits of SG and LG applications on coriander plant health and growth.

### The Influence of Amino Acids on Chloroplast Density in Parsley Stem Cells

Using optical microscopy at 400x magnification, the chloroplast densities within parsley stem base skin cells were examined. Glass carriers were employed to prepare the slides, ensuring a consistent and clear view of the cellular structures. The investigation was structured to include a control group (Figure 3A) and two experimental groups treated with seaweed amino acid (Figure 3B) and L-amino acid (Figure 3C) at 100 ppm concentrations, applied every three days over a period of nearly two months, from January 15 to March 9, 2024. Post-treatment, the cells were imaged to ascertain the differential effects of the amino acid applications on the quantity and distribution of chloroplasts, which are pivotal to photosynthetic efficiency. The resulting images, including a diagrammatic representation (Figure 3D), provided a qualitative visual representation of the changes induced by the treatments, laying the groundwork for further quantitative analysis.

### Data Collection and Analysis

On the date of harvest, March 9, 2024, data collection was carried out to compare the growth outcomes of the different treatment groups. The primary focus was on the qualitative growth aspects, such as the density of foliage, and a quantitative measure— plant height. For the qualitative assessment, visual inspection of foliage was performed, noting the general health and vigor of the plants. Plant height was measured from the base of the stem to the apex of the plant using a standard ruler, with care taken to ensure accuracy and consistency across all samples. The recorded heights provided the data for a comparative analysis between the treated and control groups. The implications of these results were considered in the context of amino acid application as a potential agronomic practice for enhancing crop yield and quality.

## Results

### The Efficacy of Amino Acid Treatments on Coriander

The experimental results depicted in Figure 1 provide a stark visual contrast between coriander plants subjected to different treatments. The plants treated with Seaweed amino acid (SG) and L-amino acid (LG) at a concentration of 100 ppm exhibit a pronounced increase in biomass compared to the control group (CG), which received no amino acid application. This enhancement is particularly evident in the density of the foliage, which suggests an improved nutritional status of the plants receiving amino acid applications. The root application method ensured direct availability of the amino acids to the plant roots, potentially facilitating a more efficient uptake than foliar applications.

**Figure 1:**
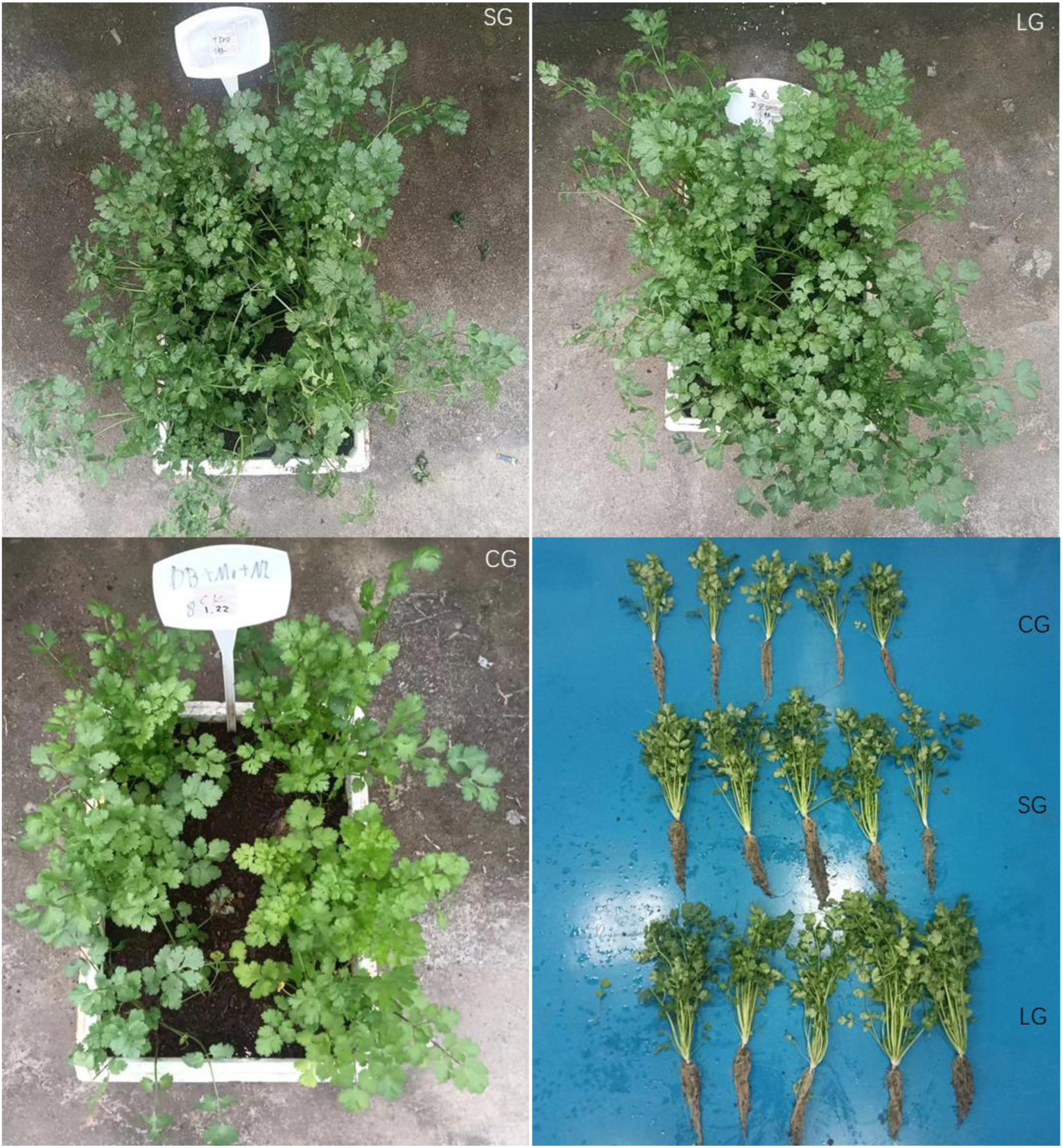
Comparative Analysis of Coriander Growth Response to Root Applications of Seaweed Amino Acid (SG) and L-amino Acid (LG). Top Left (SG): Coriander plants treated with seaweed amino acid at 100 ppm, showing dense foliage. Top Right (LG): Coriander plants treated with L-amino acid at 100 ppm, exhibiting similar vegetative growth. Bottom Left (CG): Control coriander plants irrigated only with water, with noticeably less growth. Bottom Right: Comparative height analysis of harvested coriander plants, demonstrating the impact of SG and LG treatments versus the control group.

A closer examination reveals that both SG and LG treatments result in plants with more vigorous growth than the control group, as indicated by the lush, green appearance of the treated plants’ leaves. This is consistent with the known role of amino acids in plant physiology, where they serve as building blocks for proteins and influence various growth-related processes such as chlorophyll synthesis and stress response. Moreover, the similarity in growth patterns between the SG and LG groups suggests that both types of amino acid applications can be effective in promoting coriander growth under the conditions provided in Guangzhou, China.

The fourth subfigure provides a comparative analysis of plant height, a quantitative measure of growth. The height metrics align with the qualitative observations of foliage density, confirming the beneficial effects of SG and LG treatments. The fact that the control group plants are noticeably shorter and less robust underscores the potential of amino acid applications as a means to enhance crop yield and quality. It is also indicative of the sufficiency of 100 ppm concentration for noticeable effects on coriander growth, although further research could explore the optimization of dosage and application frequency for maximal yield improvement.

### Root Development Responses to Amino Acid Applications in Coriander

The comprehensive analysis of coriander root development under Seaweed amino acid (SG) and L-amino acid (LG) treatments provides valuable insights into the subterranean effects of these biostimulants. In the presented Figure 2, the root systems of the coriander plants treated with SG and LG exhibit a pronounced improvement in development over the control group (CG). This improvement is not limited to length but includes the overall root biomass and architecture, which are critical factors for nutrient and water uptake. The robustness of the root structures in the SG and LG groups could also indicate enhanced resilience to environmental stressors, leading to a more sustainable growth cycle. Such root morphology is likely the result of improved metabolic processes fostered by the amino acid treatments, which can increase the activity of root-expanding enzymes and stimulate growth-promoting hormones.

**Figure 2.**
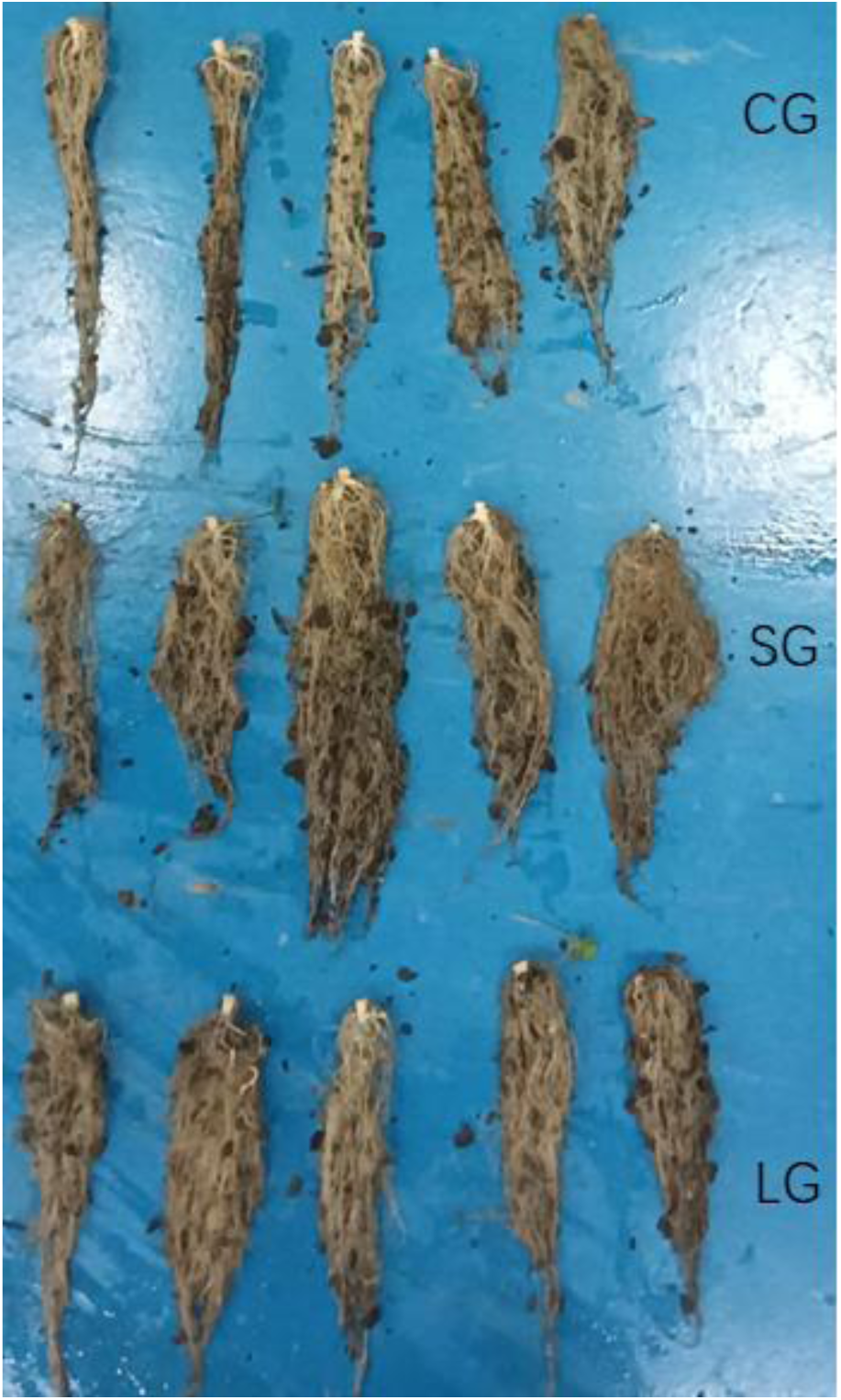
Differential Impact of Seaweed Amino Acid (SG) and L-amino Acid (LG) Treatments on Coriander Root Morphology. The image presents a comparative view of coriander root systems after treatment with Seaweed amino acids (SG) and L-amino acids (LG) against a control group (CG). The control group roots (top row) display a baseline development, having been irrigated solely with water. The SG group roots (middle row) exhibit enhanced development, indicative of the positive influence of Seaweed amino acid treatment. In contrast, the LG group roots (bottom row) show a comparable advancement in root structure to the SG group, suggesting both amino acid treatments contribute to improved root architecture and possibly, plant resilience at harvest.

Comparing the root systems of the SG and LG treatments, there is a notable similarity in root growth enhancement, suggesting that both treatments have a comparable beneficial effect on the plants’ root architecture. However, without quantitative data such as root length, surface area, and volume measurements, it is challenging to draw definitive conclusions about the superiority of one treatment over the other. The uniformity in the enhanced root development among the SG and LG groups implies that the application of amino acids, regardless of their source, could be a crucial agronomic practice to boost the growth of crops like coriander. The integration of such biostimulants into conventional farming practices could potentially lead to increased crop yields, particularly in regions where soil fertility and water availability may be limiting factors.

### Parsley Height Response to Amino Acid Treatments

Table 1 presents a comparative analysis of parsley height following treatments with seaweed amino acid, L-amino acid, and a control group. Parsley treated with L-amino acid demonstrated the highest average height at 47.56 cm, suggesting a significant positive response compared to the control group, which had the lowest average height at 37.76 cm. The seaweed amino acid treatment resulted in a moderately high average height of 45.94 cm, indicating its effectiveness, though slightly less so than the L-amino acid. The control group’s significantly lower average height underscores the efficacy of amino acid treatments in promoting plant growth. The variance in growth response among treatments highlights the potential of specific amino acids in enhancing plant height, with L-amino acid showing the most substantial impact.

**Table 1.** The height of parsley.

| Treatments | Height (cm) |  |  |  |  | Average |
| --- | --- | --- | --- | --- | --- | --- |
| Seaweed amino acid | 42.5 | 50.3 | 45.7 | 44.7 | 46.5 | 45.94 |
| L-amino acid | 47.2 | 48.3 | 44.7 | 49.8 | 47.8 | 47.56 |
| Control | 43.4 | 33.1 | 33.2 | 44.1 | 35 | 37.76 |

### Parsley Stem Width Response to Different Treatments

The data presented in Table 2 indicates that parsley plants treated with seaweed amino acids exhibited the highest average stem width at 20.68 mm, suggesting a positive impact on stem growth compared to the other treatments. The L-amino acid treatment yielded a slightly lower average stem width of 19.6 mm, still above the control group, which had the lowest average stem width of 14.78 mm. This variation in stem width across the treatments suggests that the type of amino acid used can significantly influence the physical growth attributes of parsley, with seaweed amino acids showing the most considerable enhancement in stem thickness. The control group’s results, which represent normal growing conditions without added treatments, establish a baseline that highlights the effectiveness of amino acid-based treatments in promoting stem growth.

**Table 2:**
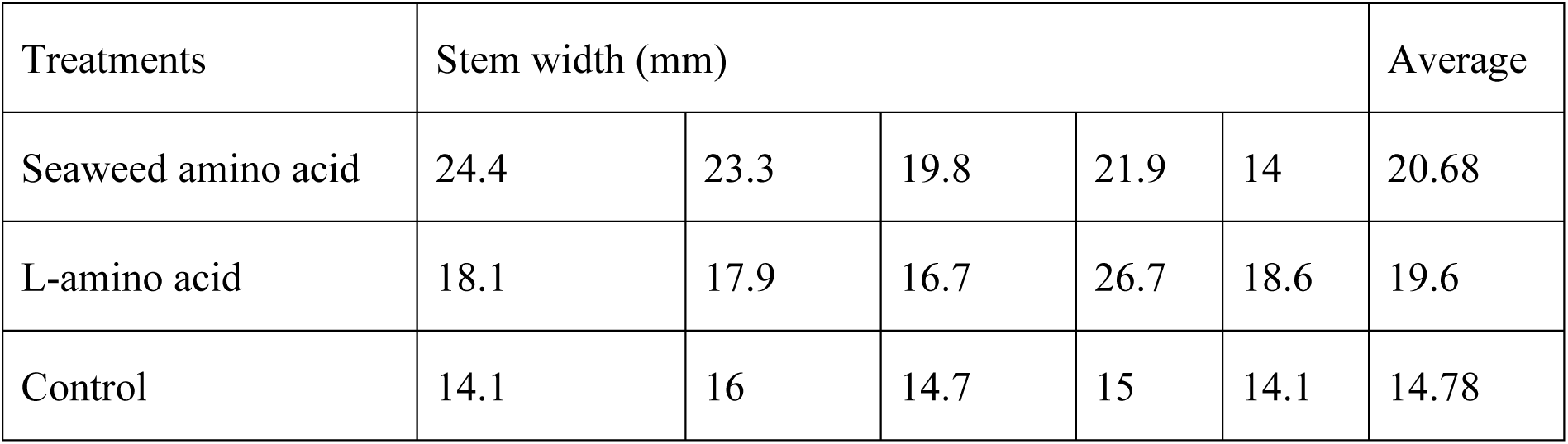
The stem width of parsley.

| Treatments | Stem width (mm) |  |  |  |  | Average |
| --- | --- | --- | --- | --- | --- | --- |
| Seaweed amino acid | 24.4 | 23.3 | 19.8 | 21.9 | 14 | 20.68 |
| L-amino acid | 18.1 | 17.9 | 16.7 | 26.7 | 18.6 | 19.6 |
| Control | 14.1 | 16 | 14.7 | 15 | 14.1 | 14.78 |

### The Impact of Amino Acid Treatments on Parsley Chlorophyll Content

The data presented in Table 3 illustrates the effect of different amino acid treatments on the chlorophyll content of parsley, measured in SPAD units. Specifically, parsley treated with seaweed amino acid exhibited an average chlorophyll content of 61.92 SPAD, indicating a positive impact on chlorophyll synthesis or preservation. The treatment with L-amino acids showed an even more pronounced effect, yielding the highest average chlorophyll content of 68.32 SPAD, suggesting that this specific treatment might be more effective in enhancing chlorophyll content in parsley. In contrast, the control group, which presumably did not receive any amino acid treatment, had a significantly lower average chlorophyll content of 50.66 SPAD. These results suggest that amino acid treatments, particularly L-amino acids, can significantly enhance the chlorophyll content in parsley, potentially indicating improved health or growth conditions for the plants treated with these substances.

**Table 3:** The contents of parsley chlorophyll.

| Treatments | Chlorophyll content (SPAD) |  |  |  |  | Average |
| --- | --- | --- | --- | --- | --- | --- |
| Seaweed amino acid | 56.3 | 63.6 | 56.5 | 66.2 | 67 | 61.92 |
| L-amino acid | 60.7 | 73.8 | 65.9 | 67.5 | 73.7 | 68.32 |
| Control | 57.8 | 58.1 | 45.4 | 42.7 | 49.3 | 50.66 |

### Comparative Analysis of Chloroplast Density (Figure 3A, 3B, 3C)

The visual assessment of Figure 3A (Control), Figure 3B (Seaweed amino acid), and Figure 3C (Left-handed amino acid) reveals a distinct pattern in chloroplast concentration across the experimental groups. The control group (Figure 3A) presents with a basal chloroplast dispersion characteristic of a standard photosynthetic profile within the parsley stem cells—highlighting a relatively sparse and evenly distributed chloroplast population. Contrastingly, the cells treated with seaweed amino acid (Figure 3B) exhibit a notable uptick in chloroplast density, marked by a pronounced green vibrancy. This suggests an augmented photosynthetic capacity or a cellular adaptation to the seaweed-based nutrients. The most profound effect is seen in the L-amino acid-treated cells (Figure 3C), where the chloroplasts’ density peaks, signified by tightly packed clusters and heightened green color intensity. This dense accumulation likely signifies a substantial bolstering of the photosynthetic apparatus, possibly driven by the specific biological activity of the applied stereoisomer.

**Figure 3.**
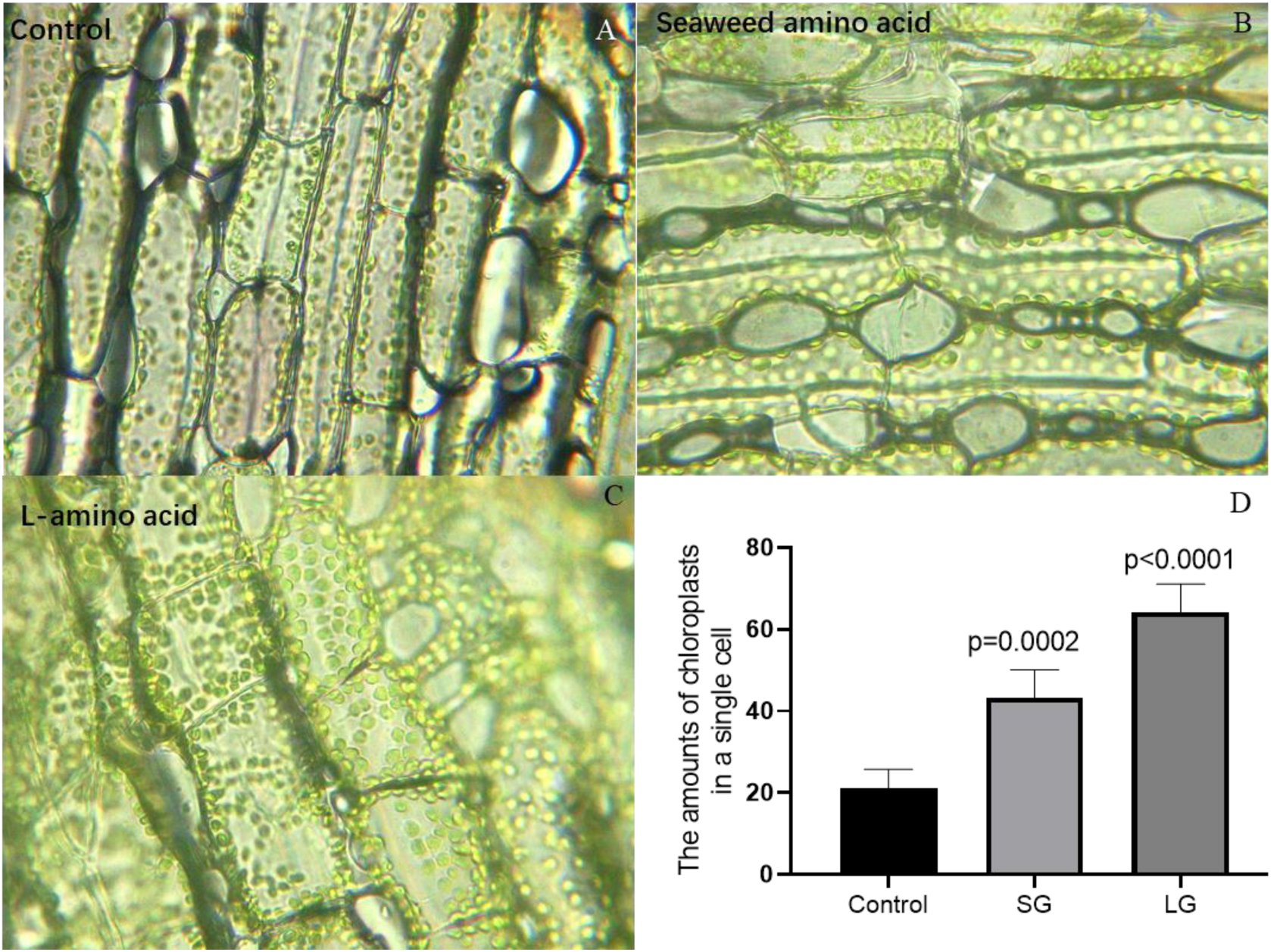
Comparative Analysis of Chloroplast Density in Parsley Stem Cells Under Different Treatments.Panel A (Control) depicts parsley stem cells with a baseline chloroplast density. Panel B (Seaweed Amino Acid) shows cells treated with 100 ppm seaweed amino acid, demonstrating an increased chloroplast density. Panel C (L-Amino Acid) exhibits cells treated with 100 ppm L-amino acid, showing the highest density of chloroplasts among the groups. The graphical inset (bottom right) provides a visual summary of the chloroplast density, with each bar representing the average chloroplast count per cell from five random cells in each treatment group. Chloroplast counts were estimated through manual inspection of micrographs taken between January 15 and March 9, 2024 at Zengcheng, Guangzhou, China.

### Implications of Amino Acid Treatment on Chloroplasts (Figure 3D)

The implications drawn from Figures 3A, 3B, and 3C are twofold. First, the increase in chloroplast content post amino acid treatments, particularly with the L-variant (Figure 3C), suggests that amino acids can act as significant modulators of plant cell physiology, enhancing chloroplast numbers and/or functionality. Such an effect aligns with the role of amino acids as biostimulants, modulating plant growth and stress response pathways. Second, the pronounced response to L-amino acid treatment underscores a potential chirality-dependent mechanism at play within the parsley stem cells (Figure 3D), hinting at sophisticated cellular recognition pathways that may be harnessed for precision agriculture to maximize photosynthetic efficiency. Further investigations, however, should involve a broader sample population, tightly controlled experimental conditions, and rigorous quantitative methodologies to validate these initial visual interpretations and unravel the complex biochemistry involved.

## Discussion

The observed enhancement in coriander growth due to the application of Seaweed amino acid (SG) and L-amino acid (LG) treatments has significant agricultural implications. The robust vegetative growth in the SG and LG groups, compared to the control (CG), indicates a promising potential for these treatments to serve as bio-stimulants. The application of amino acids can lead to improved nutrient uptake, stress tolerance, and overall plant vigor, which are essential attributes for high-yield agriculture. These findings align with previous research suggesting that amino acids can stimulate root development, thereby facilitating a greater absorption of water and nutrients from the soil ^14^. Moreover, this increased vegetative mass could also contribute to soil health through a higher rate of organic matter return to the soil post-harvest.

The similarity in growth responses to SG and LG applications raises questions about the relative efficacy of these two types of amino acids. While both have shown positive effects on the coriander plants’ growth, it remains unclear if one is superior or if they are equally beneficial in different conditions. The lack of significant visible difference between the two treated groups in this experiment suggests that the specific type of amino acid might not be as crucial as the presence of amino acids in general. However, further research could elucidate if there are any long-term differences in plant health or if there are specific circumstances under which one type outperforms the other ^15^.

Before recommending SG and LG amino acid treatments for widespread agricultural use, several factors must be considered. The cost-effectiveness of these treatments needs to be evaluated, especially since the economic feasibility is as critical as the biological efficacy for adoption by farmers. Additionally, the environmental impacts of scaling up amino acid applications should be assessed. While the treatments are derived from natural sources, their production, transportation, and application at large scales could have unforeseen ecological consequences ^16^. Further studies should also investigate the optimum concentration and application frequency to maximize the benefits of amino acid treatments while minimizing any negative effects. The possibility of integrating these treatments into a holistic plant care regimen, including traditional fertilization and pest management strategies, could pave the way for more sustainable and productive agricultural practices.

The discernible differences in root morphology between the coriander plants treated with Seaweed amino acid (SG), L-amino acid (LG), and the control group (CG) underscore the role of specific amino acid applications in root system development. The SG and LG treatments appear to foster more robust root growth compared to the control, suggesting that these amino acids may facilitate better nutrient and water uptake, potentially leading to improved overall plant health and yield. The resemblance between the SG and LG treated plants indicates that both types of amino acids can be similarly beneficial for root development. However, further research is required to quantify these observations and understand the underlying mechanisms, ensuring that such treatments can be optimized and reliably integrated into agricultural practices for enhanced crop production.

The results from the study on parsley growth response to amino acid treatments provide compelling evidence on the role of amino acids in plant development, particularly in terms of plant height and stem width. The application of L-amino acids led to a notable increase in plant height, surpassing both the control group and the group treated with seaweed amino acids. This suggests that L-amino acids may possess unique properties or mechanisms that stimulate vertical growth in parsley more effectively than seaweed-derived amino acids or natural growth conditions without supplementation. The significant difference in height between the treated groups and the control underscores the potential of amino acid treatments as a means to enhance plant growth, which could have implications for agricultural practices aimed at increasing crop yield and optimizing space usage through vertical growth^17, 18^.

In contrast, the stem width response to different amino acid treatments reveals a different aspect of plant growth influenced by amino acid composition. Seaweed amino acids resulted in the greatest increase in stem width, suggesting that these amino acids may support broader developmental processes within the plant, contributing to a sturdier and potentially more resilient growth form^19^. The disparity between the effects of L- and seaweed amino acids on stem width versus height indicates that the biochemical pathways influenced by these treatments may differ, with specific amino acids triggering distinct growth responses. This variation highlights the complexity of plant growth and the potential for targeted growth modulation through the application of different types of amino acids^20^.

These findings open new avenues for research into the mechanisms behind the growth-promoting effects of amino acids in plants. Understanding how different amino acids influence growth parameters such as height and stem width could lead to the development of optimized treatment regimens that leverage the unique benefits of both L- and seaweed amino acids. For agricultural and horticultural practices, such insights could translate into strategies that enhance plant resilience and productivity through the targeted application of amino acid treatments. Additionally, the pronounced growth responses observed in this study underscore the potential of amino acid treatments as a sustainable and natural alternative to synthetic growth promoters, aligning with the increasing demand for organic farming methods.

The study on the impact of amino acid treatments on parsley highlights intriguing insights into the potential of these treatments to enhance chlorophyll content and, by extension, the health and growth efficiency of plants. The observed increase in chlorophyll levels, particularly with the application of L-amino acids, points towards an efficacious strategy for boosting photosynthetic capacity. Chlorophyll is fundamental to photosynthesis, the process through which plants convert light energy into chemical energy. An increase in chlorophyll content, as indicated by higher SPAD units, suggests an enhanced ability of plants to absorb light, thereby potentially improving their growth rates and health. The stark contrast between the treated groups and the control underscores the significance of amino acid treatments in agricultural practices, especially in optimizing plant nutrition and maximizing growth outcomes.

The comparative analysis of chloroplast density across different treatment groups further elucidates the beneficial effects of amino acid treatments on the microstructural level. Chloroplasts, being the sites of photosynthesis, play a pivotal role in plant vitality^21^. The visual evidence from the study showing increased chloroplast density in treatments, especially with L-amino acids, suggests not just an increase in chlorophyll content but possibly an enhanced structural capacity for photosynthesis^22^. This augmented density and the associated vibrancy in green color could translate into a higher photosynthetic efficiency, allowing plants to convert light energy into chemical energy more effectively. The differentiation in response between the seaweed amino acid and L-amino acid treatments also hints at the specificity of plant responses to different types of amino acids, suggesting a potential avenue for tailored nutritional strategies to optimize plant photosynthesis and growth^23, 24^.

Finally, the implications of these amino acid treatments on chloroplast content and functionality spotlight the role of amino acids as potent biostimulants that can modulate plant physiology. The particularly pronounced response to L-amino acids could indicate a chirality-specific interaction within plant cells, revealing an intricate layer of biochemical specificity that plants may utilize to discern and respond to different molecular configurations. Such findings not only deepen our understanding of plant biology and physiology but also open up new possibilities for the development of precision agriculture techniques. By harnessing the specific effects of amino acid chirality, agricultural practices could be refined to target and enhance desired plant responses, such as photosynthetic efficiency, thereby contributing to sustainable and efficient crop production^25^. Future research should aim to unravel the underlying mechanisms of these chirality-dependent responses and explore their practical applications in agriculture.

In conclusion, this study contributes valuable insights into the role of Seaweed in promoting coriander growth and enhancing its anti-stress ability. The differential impacts of foliar and root applications underscore the importance of tailored agronomic practices in leveraging the benefits of biostimulants. Moreover, the findings advocate for the integration of biostimulants like Seaweed into sustainable agricultural strategies, offering a promising avenue for improving crop resilience and yield in the face of escalating environmental stresses. Further research is warranted to explore the mechanistic basis of Seaweed action in plants and to extend these observations to field conditions, where the interplay of biostimulants with diverse environmental factors can be more comprehensively assessed.

